# Reduced food accessibility enhances value-based decision-making in rats

**DOI:** 10.64898/2026.10.20.759714

**Authors:** Clara I. González Matellán, Vanesa Hurtado-Nieves, F. Javier Aguado, Marta Bourdelande García, María A. Bermúdez, Jose L. Pardo-Vazquez

## Abstract

Finding nutrients regularly is critical for survival; therefore, most animals invest a large portion of their time and energy in securing access to food. Food acquisition activates the brain’s reward system, enhancing motivation and thereby increasing chances to succeed again in the future. However, little is known about the effect of food accessibility on cognition. We hypothesize that reducing food accessibility by requiring laboratory animals to get it with effort, will expose them to daily learning challenges, enhance reward system engagement and consequently have a positive impact on cognitive performance. To test whether effortful access to food confers cognitive benefits, one group of rats was housed with a food dispenser specifically designed to reduce food accessibility (Low Accessibility, LA). The comparison group had free access to food at any time (High Accessibility, HA). Both groups were trained in a value-based task in which they were required to choose between two options and, through trial and error, learn which option was more likely to deliver a reward. We found that both groups learned the task structure similarly. However, the LA group showed a higher proportion of optimal choices, indicating better task performance. We also found that LA rats were more sensitive to the outcomes of previous decisions. Collectively, our findings suggest that reducing food accessibility improves value-based decision-making, potentially by increasing sensitivity to action outcomes, highlighting a previously underappreciated link between food acquisition demands and cognitive performance.

## Introduction

Humans evolved under ecological conditions in which food resources were often scarce and obtaining them typically required physical and cognitive effort. Consequently, natural selection favored highly efficient physiological and behavioral mechanisms that promote food seeking and consumption, whereas mechanisms limiting energy intake are comparatively slower and weaker. In modern environments characterized by the widespread availability of highly palatable, energy-dense foods, this evolutionary mismatch is thought to contribute substantially to the high prevalence of overweight and obesity (Hill & Peters, 1998; Karaoglan & Karaoglan, 2024; Speakman, 2013; Speakman & Elmquist, 2022). A related concern applies to animal research. Because securing nutrients is essential for survival, free-living animals devote a considerable proportion of their time and energy to foraging and other behaviors that ensure future access to food. These activities continuously engage cognitive processes involved in learning, behavioral adaptation, and decision-making, ultimately increasing the likelihood of survival and reproductive success. Consistent with contemporary theories of reinforcement learning, successful foraging depends on an animal’s ability to learn relationships between actions and their consequences and to update behavior according to changing environmental contingencies (Gershman & Uchida, 2019; Sutton & Barto, 2018). Laboratory rodents maintained under standard *ad libitum* feeding conditions differ markedly from their wild counterparts in several physiological and behavioral traits, raising questions about the ecological validity and translational relevance of some experimental models (Feige-Diller et al., 2020; Hinde et al., 2018; Keenan et al., 1996; Martin et al., 2010).

Two essential dimensions of feeding conditions are food availability and food accessibility. Food availability refers to the quantity of food that is available to an organism, whereas food accessibility concerns the extent to which obtaining that food requires cognitive and/or physical effort. The consequences of reduced food availability have been extensively investigated through dietary interventions such as caloric restriction and intermittent fasting. These interventions have been associated with improvements in healthspan and lifespan across a range of animal species and produce favorable effects on metabolic and physiological markers in humans (Ching & Hsu, 2025; Green et al., 2022; Mattison et al., 2017; Nakagawa et al., 2012; Speakman, 2020). Furthermore, a substantial body of evidence indicates that dietary restriction can enhance cognitive functions, including learning and memory, and may contribute to improved brain health (Kuhla et al., 2013; Lobo et al., 2022; Lü et al., 2023; Ma et al., 2018; Pani, 2015; Singh et al., 2012; Witte et al., 2009). Taken together, these findings suggest that reducing food availability can counteract some of the detrimental consequences of overweight and obesity. Despite the extensive research into dietary restriction, the effects of food accessibility on cognition and health remain poorly understood. Investigating how the effort required to obtain food influences behaviour and brain function may therefore provide new insights into how feeding conditions shape cognitive processes. In addition, introducing feeding regimens that more closely resemble natural conditions may improve the ecological validity and translational relevance of animal models used in biomedical research (Feige-Diller et al., 2020; François et al., 2022; Graham, 2021).

Research examining the relationships between feeding conditions and cognition still presents important limitations. First, food accessibility, that is, whether animals must exert effort to obtain food, has been mostly ignored. Second, many experimental tasks are directly related to foraging, making it difficult to determine whether the observed effects generalize beyond food-related contexts. Finally, most studies have relied on relatively simple behavioral paradigms, and consequently little is known about how dietary manipulations affect higher-order executive functions such as decision-making.

Decision-making is a high-level cognitive process through which an organism selects a course of action from multiple alternatives, typically with the goal of maximizing benefits or minimizing adverse outcomes (Morelli et al., 2022). Animals make countless decisions every day, many of which have profound consequences for survival and reproduction (Wispinski et al., 2020). Importantly, deficits in decision-making have been identified across numerous neuropsychiatric and neurological disorders, including substance use disorders, depression, bipolar disorder, and Parkinson’s disease (Adida et al., 2011; Ariesen et al., 2023; Chen et al., 2020; Colautti et al., 2021; Gu et al., 2020; Perugini et al., 2018; Rahman et al., 2001; Verdejo-García et al., 2019). Consequently, understanding the behavioral and neural mechanisms underlying decision-making has long been a central objective of behavioral and cognitive neuroscience (Hauber, 2026; Rahman et al., 2001; Winstanley & Floresco, 2016).

Previous research has demonstrated that food availability influences cognitive performance in both humans and rodents, particularly in domains such as learning and memory. However, beyond the quantity of food available, the way food is obtained may also be a critical determinant of higher-order cognition. We hypothesize that food accessibility modulates cognitive and neural processes involved in adaptive decision-making. Therefore, providing laboratory animals with a more meaningful and biologically relevant interaction with their environment—by requiring them to expend effort to obtain food—will expose them to learning opportunities on a daily basis and enhance the efficiency of reward-related neural circuits even in situations unrelated to food procurement. This work has the potential to advance the development of more ecologically relevant animal models for behavioral neuroscience and may ultimately inform the study of neuropsychiatric disorders, in which decision-making deficits represent a prominent and clinically significant feature.

## Materials and Methods

### Animals

Twelve female Long-Evans rats (Janvier Labs) were used in this study. Animals were 35 days old at the beginning of the experiment. They were cared for according to the guidelines of the Physiological Spanish Society, International Council for Laboratory Animals Science, and the European Union guidelines (2010/63/EU), and the experimental protocol was reviewed and approved by the Ethics Committee for animal research of the Universidade da Coruña and Xunta de Galicia (ref. 15012/2022/010).

### Experimental design

Rats were randomly assigned to one of two experimental groups: High Accessibility (HA) or Low Accessibility (LA). They were housed in pairs, with each cage containing one HA and one LA animal. Cages were equipped with a custom-made transparent divider that created two independent compartments while allowing social interaction through multiple perforations distributed along the separator. Animals were maintained under controlled temperature and humidity conditions on a 12 h light/dark cycle, and all experimental procedures were conducted during the light phase.

Following arrival, rats underwent a 14-day habituation period during which they had *ad libitum* access to food and water. Food intake and body weight were monitored at least 4 times per week (Figure 1A). Subsequently, animals assigned to the LA group were exposed to a custom-designed feeding system intended to reduce food accessibility by making animals get the food with effort. This device required rats to open a series of gates positioned at different heights to gain access to food pellets. The feeder was introduced gradually over the course of one week. On the first day, pellets were placed in the lowest compartment. Each subsequent day, the pellets were moved one level higher. By the end of this habituation period, all rats were able to open every gate and reliably obtain food from the highest compartment. In contrast, HA animals retained unrestricted access to food throughout the experiment. These housing and feeding conditions were maintained for 6 weeks before the start of behavioral training and remained unchanged throughout the entire experiment.

**Figure 1.**
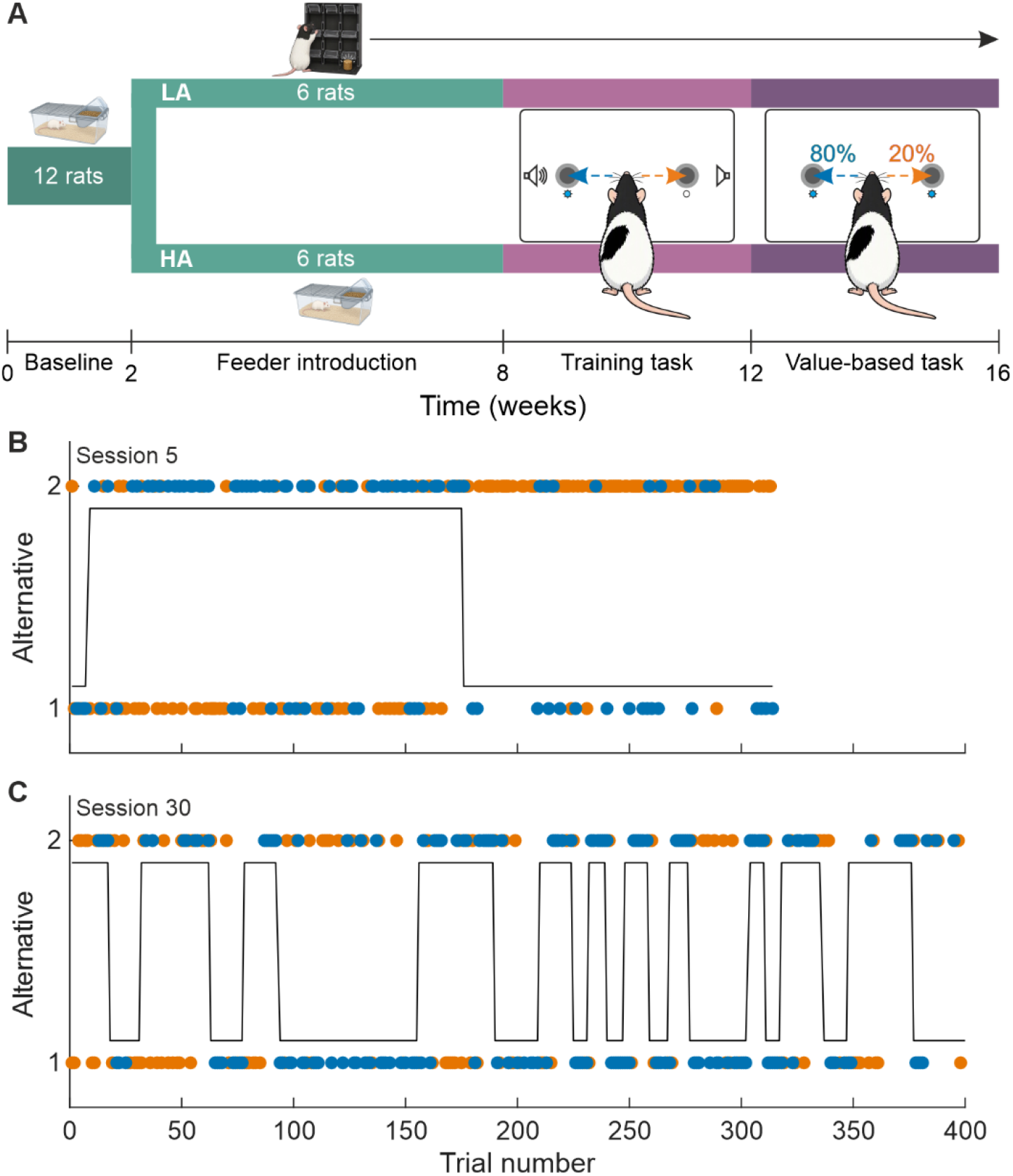
Experimental design and performance in the value-based task. **A**, Schematic representation of the experiment’s timeline; LA, low food accessibility, HA, high food accessibility. **B**, **C**, Trial-by-trial performance of a representative rat during an early (B) and a late (C) session of the value-based task. Each dot represents the rat’s choice on a given trial (alternative 1 or alternative 2). Blue dots indicate rewarded trials, whereas orange dots indicate non-rewarded trials. The solid black line denotes the optimal alternative, which switched whenever the rat selected the optimal option on six consecutive trials.

Behavioral training was conducted in custom-made operant cages. During training, animals were subjected to a water-restriction schedule in which water was obtained primarily through task performance. Correct responses were rewarded with a water drop delivered through a solenoid valve (The Lee Company, LHDA0531115H) connected to a reservoir. Animals received *ad libitum* access to water during Friday afternoons and Saturdays, and supplemental water was provided when necessary to maintain body weight above the predetermined welfare threshold (≥ 85% of baseline body weight). HA and LA animals were trained in parallel in independent behavioral cages, and cage assignment was systematically rotated across sessions to minimize potential apparatus-related biases.

Animals were initially trained to perform a sequence of actions (poke in the central port – poke in a lateral port – poke in the reward port) using sound and light to guide them (training task, TT). After 23 sessions, the animals were transitioned to a value-based decision-making task (VT), from which 30 sessions were collected for analysis. Each session in the TT and VT lasted for 90 and 70 minutes, respectively.

### Behavioral tasks

#### Training task (TT)

Behavioral testing was conducted in square operant cages housed within sound-attenuating cubicles. Each chamber was equipped with four nose-poke ports. Two ports were positioned on the central wall: an upper port was used to initiate trials and a lower port for reward delivery. The remaining two ports, located on the left and right sides of the chamber beneath a speaker (Tweeter XPL model XTW2505), served as the response options. All nose-poke ports were equipped with infrared detectors for response detection and integrated LED lights.

The trial could be initiated when the LED above the central start port was illuminated, indicating its availability. A trial began when the animal inserted its snout into this port. Following trial initiation, a compound audiovisual cue consisting of an illuminated LED and a burst of white noise was presented on either the left or right side of the chamber. Rats were required to identify the location of the cue and respond by nose-poking the corresponding lateral response port.

Correct responses triggered illumination of the reward port and delivery of a 30-µL water reward upon entry into the port. Incorrect responses resulted in a 5 s timeout period, after which a new trial could be initiated. Animals were allowed a maximum of 20 s to make each required response. Failure to respond within this time window resulted in trial termination and a 1.5 s timeout before the start of the next trial. The intertrial interval was set to 1.5 s.

#### Value-based decision-making task (VT)

Testing was conducted in V-shaped operant cages equipped with four nose-poke ports arranged as follows: a central initiation port, a central reward port, and two response ports in the opposite wall. All ports were equipped with infrared detectors and cue LEDs. Trial initiation, response requirements, time limits, and reward collection were identical to those used in the TT.

Trials were self-initiated by a nose poke into the illuminated central port. Following trial initiation, rats were required to choose between the left and right response ports in the absence of external differential sensory cues (both ports were illuminated to indicate availability). One option, the optimal, was associated with an 80% probability of reward, whereas the alternative option was rewarded on 20% of trials (the initial assignment of reward probabilities was randomized). Rewarded choices triggered illumination of the reward port, where a 30-µL water reward could be obtained. Non-rewarded choices were followed by a 5-second timeout penalty. Animals were therefore required to identify the current optimal option through trial-and-error learning.

The reward contingencies remained stable until the animal made six consecutive choices of the optimal option, at which point the reward probabilities were reversed, and a new block began. Thus, successful task performance required continuous adaptation to changing reward contingencies (Figure 1B and C).

### Data analysis

We analyzed several behavioral variables to fully characterize performance. All data analyses were performed using R [v. 4.3.1; (R Core Team, 2023)] with the packages lme4 [v. 1.1.34; (Bates et al., 2015)], lmerTest [v. 3.2.1; (Kuznetsova et al., 2017)], glmmTMB [v. 1.1.14; (Brooks et al., 2017)] and emmeans [v. 1.9.0; (Lenth & Piaskowski, 2026)].

Body weight and food intake were analyzed separately for each experimental phase (baseline, food accessibility, TT and VT). Body weight was analyzed using linear mixed-effects models (LMMs) fitted by maximum likelihood, with accessibility group, days (centered at the midpoint of each phase) and their interaction as fixed effects, weighing day (Monday vs. other days) as a covariate, since body weight was higher after *ad libitum* water access during weekends, and subject identity as a random intercept. Body weight at arrival and at the end of training was compared between groups using Welch’s t-test. Daily food intake was computed from the amount of food consumed between consecutive records and analyzed using an LMM with accessibility group, phase and their interaction as fixed effects, interval type (with or without a weekend of *ad libitum* water access) as a covariate, and subject identity as a random intercept. Group differences within each phase were estimated using contrasts of estimated marginal means.

On the one hand, count variables were analyzed using generalized linear mixed-effects models (GLMMs). For completed and optimal trials, a binomial error distribution with a logit link function was used, while for number of completed blocks we used a Poisson distribution with a log link function. Fixed effects included accessibility group (LA vs. HA), session number, and their interaction. Subject identity was included as a random intercept. When we observed a substantial overdispersion, an observation-level random effect (OLRE) was added to model extra-binomial variation (Harrison, 2015). Model parameters were estimated by maximum likelihood using the Laplace approximation.

The cumulative number of blocks, on the other hand, was analyzed by fitting a linear mixed-effects model (LMM) using restricted maximum likelihood (REML). Accessibility group and session (mean-centered) were included as fixed effects, together with their interaction. To account for the repeated-measures structure of the data, a random intercept was included for each animal (subject identity). Statistical significance of fixed effects was assessed using t-tests with Satterthwaite approximations for the degrees of freedom.

To assess the influence of previous outcomes on choice behavior, we analyzed trial-by-trial stay decisions using a generalized linear mixed-effects model (GLMM) with a binomial error distribution and logit link function. The dependent variable (stay) indicated whether the animal repeated the same choice made on the previous trial. Fixed effects included accessibility group (LA vs. HA), reward outcomes received on the previous four trials, session number (mean-centered), and the interactions between group and each reward-history term. Rat identity was included as a random intercept to account for repeated observations within subjects.

## Results

We first examined whether manipulating food accessibility altered food intake and body weight throughout the experiment. During baseline, the two groups did not differ in either food consumption or body weight. HA and LA rats consumed 13.7 ± 0.6 and 14.0 ± 0.6 g/day, respectively (*β* = 0.36 ± 0.60 g/day, *p* = 0.55), and weighed 108.7 ± 3.5 g and 107.8 ± 4.4 g, respectively (*p* = 0.73). Following feeder introduction and before training-task sessions began, LA rats consumed 11% less food than HA rats (12.9 ± 0.7 vs. 14.6 ± 0.9 g/day; *β* = −1.73 ± 0.47 g/day, *p* = 0.001) and gained weight at a significantly slower rate (0.38 g/day less, *p* < 0.001). During the TT phase, LA rats showed a modest but significant reduction in food intake relative to HA rats (*β* = −1.37 ± 0.50 g/day, *p* = 0.010). They also weighed, on average, 11.8 g less than HA rats, although both groups gained weight at a comparable rate (*p* = 0.69). Similarly, during the value-based task, LA rats remained significantly lighter than HA rats (*β* = −13.0 ± 5.4 g, *p* = 0.034), weighing approximately 13 g less (94.5% of HA body weight). However, no group differences were observed in either daily weight gain (*β* = −0.08 ± 0.06 g/day, *p* = 0.17) or food intake, with HA and LA rats consuming 11.5 ± 0.8 and 11.1 ± 1.0 g/day, respectively (*β* = −0.44 ± 0.48 g/day, *p* = 0.36). By the end of training, HA rats weighed 234.8 ± 12.0 g, whereas LA rats weighed 219.7 ± 10.7 g (93.5% of HA body weight, *p* = 0.043). Overall, reducing food accessibility produced a transient decrease in food intake and weight gain that resulted in persistently lower body weight in LA rats, despite the subsequent normalization of both food consumption and growth rate between groups.

To assess how animals learned the sequence of actions required to complete the VT, we first compared the proportion of completed trials (Figure 2A) using a GLMM and including an OLRE. The probability of completing a trial increased significantly across sessions (*β* = 0.020 ± 0.007 SE, *z* = 2.81, *p* = 0.005; OR = 1.02), indicating a progressive improvement in task engagement over time. In contrast, there was no evidence for a main effect of accessibility group (*β* = 0.035 ± 0.098 SE, *z* = 0.36, *p* = 0.720) and no significant accessibility group × session number interaction (*β* = −0.013 ± 0.010 SE, *z* = −1.33, *p* = 0.182), suggesting that the temporal increase in trial completion was comparable between groups. The model included 360 observations from 12 rats. The variance associated with the observation-level random effect was substantial (σ² = 0.656), confirming the presence of extra-binomial variation in the data, whereas between-subject variability was comparatively small (σ² = 0.006). Collectively, these findings indicate that animals became more likely to complete trials as training progressed, but this improvement was not influenced by experimental group.

**Figure 2.**
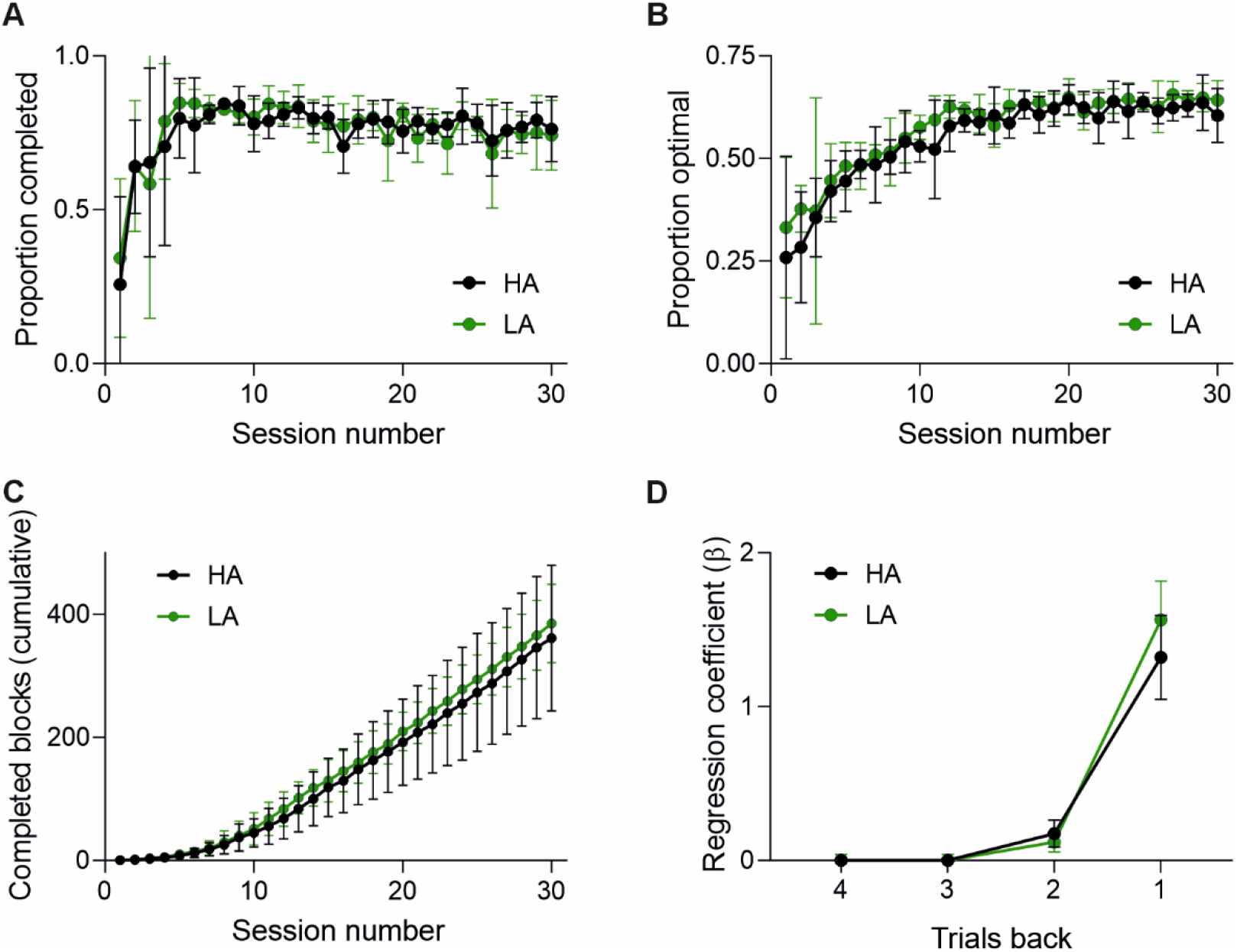
Reduced food accessibility enhances performance in the value-based task. **A**, **B**, Proportion of completed trials (A) and optimal choices (B) as a function of training session and food accessibility (HA, high food accessibility; LA, low food accessibility). **C**, Cumulative number of completed blocks as a function of training session and food accessibility; same conventions as in A and B. **D**, Regression coefficients quantifying the influence of outcomes from previous trials on current decisions. Dots and error bars represent the mean and SD across rats, respectively.

The probability of making an optimal choice increased significantly across sessions (Figure 2B), indicating progressive learning throughout training (*β* = 0.0367 ± 0.0024 SE, *z* = 15.16, *p* < 0.001). Specifically, the odds of selecting the optimal option increased by approximately 3.7% per session (OR = 1.037, 95% CI ≈ 1.032-1.042). There was also a significant main effect of accessibility group (*β* = 0.0938 ± 0.0450 SE, *z* = 2.08, *p* = 0.037), indicating that animals in the LA group exhibited a higher probability of optimal responding than those in the HA group. The estimated effect corresponds to an odds ratio of 1.10, suggesting approximately a 10% increase in the odds of making an optimal choice in the LA group. Importantly, the interaction between accessibility group and session number was not significant (*β* = −0.0032 ± 0.0034 SE, *z* = −0.94, *p* = 0.348), indicating that the rate at which performance improved across sessions did not differ between groups. Thus, both groups showed comparable learning trajectories over time, although the LA group maintained a modest overall performance advantage.

A Poisson GLMM fitted to 360 observations from 12 rats revealed a significant positive effect of session on the number of completed blocks (*β* = 0.070 ± 0.005 SE, *z* = 13.01, *p* < 0.001). This indicates that animals completed progressively more blocks as training advanced. The corresponding incidence rate ratio (IRR = exp(0.0697) = 1.072) suggests that the expected number of completed blocks increased by approximately 7.2% per session. Neither the main effect of accessibility group (*β* = 0.116 ± 0.155 SE, *z* = 0.75, *p* = 0.454) nor the accessibility group × session number interaction (*β* = −0.006 ± 0.007 SE, *z* = −0.76, *p* = 0.448) reached significance. Thus, the groups did not differ in the overall number of completed blocks, nor did they differ in the rate at which block completion increased across training. Variance partitioning indicated that unexplained variability was primarily attributable to observation-level effects (σ² = 0.218; SD = 0.467), whereas between-subject variability was comparatively modest (σ² = 0.060; SD = 0.246). This pattern suggests that most residual variation reflected session-to-session fluctuations within animals rather than stable differences between animals.

When we analyzed the cumulative number of blocks (Figure 2C), we identified a significant main effect of session number, indicating that cumulative performance increased across training sessions (*β* = 13.26 ± 0.30 SE, *t*(346) = 43.62, *p* < 0.001). In contrast, the main effect of accessibility group was not significant (*β* = 12.67 ± 23.45 SE, *t*(10) = 0.54, *p* = 0.601), suggesting that the groups did not differ significantly in performance at the mean session value. Importantly, a significant group × session interaction was observed (*β* = 0.88 ± 0.43 SE, *t*(346) = 2.05, *p* = 0.041). This interaction indicates that the rate of change in cumulative performance across sessions differed between groups. Specifically, the LA group exhibited a steeper increase in cumulative blocks over time than the HA group, consistent with a faster improvement in performance throughout training. The model was fitted to 360 observations obtained from 12 animals. The estimated SD of the random intercepts was 40.1, indicating substantial between-subject variability, whereas the residual SD was 35.3.

A binomial GLMM fitted to 111,927 observations from 12 rats revealed a strong influence of recent reward history on choice persistence (Figure 2D). The outcome of the immediately preceding trial exerted the largest effect on stay behavior (*β* = 1.330 ± 0.019 SE, *z* = 71.58, *p* < 0.001). Thus, animals were substantially more likely to repeat a choice when that choice had been rewarded on the previous trial. A smaller but still significant positive effect was observed for rewards received two trials earlier (*β* = 0.159 ± 0.019 SE, *z* = 8.59, *p* < 0.001). In contrast, reward outcomes three trials earlier exerted a modest negative influence on choice repetition (*β* = −0.053 ± 0.019 SE, *z* = −2.88, *p* = 0.004), whereas the effect of outcomes four trials back was marginal (*β* = −0.035 ± 0.018 SE, *z* = −1.91, *p* = 0.056). There was no significant main effect of accessibility group (*β* = −0.049 ± 0.080 SE, *z* = −0.62, *p* = 0.536), indicating that, in the absence of reward-history effects, overall stay behavior did not differ between groups. A significant positive effect of session number was observed (*β* = 0.023 ± 0.001 SE, *z* = 27.86, *p* < 0.001), demonstrating that animals became progressively more likely to repeat previous choices as training progressed. Critically, reward-history effects differed between groups.

The interaction between accessibility group and the most recent reward outcome was significant and positive (*β* = 0.237 ± 0.027 SE, *z* = 8.84, *p* < 0.001). This result indicates that the behavioral impact of a reward obtained on the immediately preceding trial was stronger in LA animals than in HA animals. In contrast, the interaction involving rewards received two trials earlier was negative but comparatively small (*β* = −0.058 ± 0.027 SE, *z* = −2.16, *p* = 0.031), suggesting a slightly weaker influence of second-lag rewards in the LA group. No significant group differences were observed for rewards occurring three or four trials previously (*ps* > 0.50). The variance associated with the random intercept was small (σ² = 0.017; SD = 0.130), indicating relatively limited and stable differences between animals after accounting for reward history and training effects.

## Discussion

The present study investigated whether food accessibility, independent of food availability, influences decision-making performance in rats. Animals housed under conditions that required effortful food procurement (LA) displayed a consistently higher proportion of optimal choices than animals with unrestricted food access (HA), despite showing similar rates of task acquisition, trial completion, and block completion. Furthermore, LA animals accumulated completed blocks more rapidly across training and exhibited greater sensitivity to previous outcomes. Together, these findings suggest that the way food is obtained (food accessibility), rather than merely the amount of food available, can shape cognitive processes involved in adaptive decision-making.

A central finding of this study is that reducing food accessibility improved decision accuracy without affecting overall task engagement. Both groups increased trial completion rates across sessions and completed progressively more blocks over the course of training, indicating comparable motivation to perform the task and similar acquisition of its procedural requirements. Consistent with this interpretation, the absence of an accessibility group × session interaction for optimal choices suggests that learning progressed at a similar rate in both groups, while LA animals maintained a stable performance advantage over HA animals throughout training. This pattern is theoretically important because it indicates that reduced food accessibility did not alter learning of the task structure; rather, it enhanced the quality of decision-making once the task had been acquired. Furthermore, the reward-history analysis suggests that LA rats selected the optimal option more frequently than HA rats because they were more responsive to the outcome of the preceding trial.

These findings are broadly consistent with the idea that environments requiring active resource acquisition enhance cognitive function. Enriched housing increases exposure to sensory, motor, and cognitive challenges and induces neurochemical and neuroanatomical adaptations that support cognitive performance. Experimental evidence and recent reviews indicate that enriched environments can improve learning, reduce maladaptive behaviors, and modify neural systems involved in motivation, attention, and behavioral flexibility (Domínguez-Oliva et al., 2025, 2026; Zentall, 2021). Although the feeding manipulation used here differs from conventional environmental enrichment protocols, it shares a critical feature: the animal is repeatedly exposed to situations that require active interaction with the environment to obtain biologically significant resources. The present results suggest that introducing cognitive and physical challenges into everyday feeding routines may generate some of the same benefits traditionally attributed to enrichment. However, changes in food accessibility do not merely increase environmental stimulation but can also affect physical activity, energy expenditure, and food intake.

One possible explanation is that effortful access to food repeatedly engages neural systems involved in reinforcement learning. Natural foraging requires animals to learn action-outcome contingencies, update expectations based on experience, and adapt behavior according to changing environmental conditions (Alejandro & Holroyd, 2024). Contemporary reinforcement-learning theories propose that these computations are implemented through distributed cortico-striatal circuits in which reward prediction errors drive behavioral updating. Because LA animals had to solve a sequence of problems every day to obtain food, they accumulated substantially more experience with action-outcome learning than HA animals. Over time, this repeated engagement may have strengthened cognitive processes involved in evaluating outcomes and updating decision policies (Hage et al., 2023; Mohebi et al., 2024). Such an account is consistent with our observation that LA animals showed greater sensitivity to previous outcomes. This is a behavioral signature often interpreted as more effective use of reinforcement information during decision-making.

The results are also compatible with the literature on contrafreeloading. Many species voluntarily work to obtain food even when identical food is freely available, a phenomenon that has traditionally been interpreted as reflecting the informational value of resource acquisition rather than simple caloric gain (Ferreira et al., 2021; Rosenberger et al., 2020; Rothkoff et al., 2024). Experimental and theoretical work suggests that obtaining food through effort can provide information about environmental contingencies and resource availability, thereby reducing uncertainty and improving future adaptive behavior (Frederick & Cocuzzo, 2017). From this perspective, requiring animals to actively obtain food may transform feeding from a purely consumptive activity into a continuous learning process. The superior performance of LA animals in the present task is consistent with the hypothesis that repeated engagement in cognitively demanding food procurement enhances the capacity to exploit reinforcement contingencies in contexts unrelated to feeding itself (Puppe et al., 2007; Richmond et al., 2026).

Importantly, the effects observed here cannot be readily explained by differences in food availability or caloric intake. Unlike caloric restriction and intermittent fasting paradigms, which alter energy intake and metabolic state, our manipulation targeted accessibility while maintaining food availability. Much of the previous literature demonstrating cognitive benefits of dietary interventions has focused on reduced food quantity, making it difficult to disentangle metabolic effects from behavioral effects. The present findings suggest that some cognitive advantages commonly associated with dietary restriction may instead arise, at least in part, from behavioral features of food acquisition. This distinction is important because it identifies food accessibility as a previously underappreciated variable that may contribute independently to cognitive outcomes.

From a translational perspective, these findings have implications for both laboratory animal welfare and biomedical research. Standard laboratory housing often provides continuous and effortless access to food, a condition that differs substantially from the environments under which mammalian cognitive systems evolved. Recent reviews have argued that impoverished housing conditions may influence brain function, behavior, and experimental outcomes, potentially limiting translational validity (Domínguez-Oliva et al., 2025; Hobbiesiefken et al., 2021; Kentner et al., 2021). The present results suggest that incorporating naturalistic food-procurement challenges into routine husbandry may improve ecological validity while simultaneously providing cognitive stimulation. Such manipulations could prove particularly relevant for studies of decision-making, reward processing, and neuropsychiatric disorders, where subtle differences in motivational and cognitive experience may substantially influence behavior.

Interestingly, LA rats were slightly lighter than HA rats. A stage-by-stage analysis indicated that this difference originated immediately after the introduction of the feeder, when LA rats consumed less food and consequently gained less weight than HA rats. Once established, this divergence in body weight was maintained throughout the rest of the experiment. This suggests that learning how to operate the feeder, rather than the manipulation of food accessibility itself, was the primary factor underlying the body weight difference between LA and HA rats.

Several limitations should nevertheless be acknowledged. First, although the effects of reduced food accessibility were significant, their magnitude was modest. One possible explanation is that the feeder provided only a limited cognitive and physical challenge. After learning how to operate it, within the first week, rats could access food with little effort. Future studies should test more sophisticated feeding devices that periodically modify the requirements for obtaining food, thereby maintaining a continuous learning demand. Such systems may promote greater engagement of cognitive and reward-related processes, leading to larger benefits in cognitive performance. Second, the study was limited to female rats. Considering evidence for sex-specific responses to food-related interventions (Maric et al., 2022; Oraha et al., 2022), future work should establish whether the effects of reduced food accessibility on decision-making are also present in males. Third, the sample size was modest, and although mixed-effects modelling appropriately accounted for repeated measurements, replication in larger cohorts will be important. Fourth, the present study was designed to evaluate behavioral performance and therefore cannot identify the neural mechanisms responsible for the observed effects. Direct assessment of dopaminergic signaling, cortico-striatal plasticity, or reinforcement-learning parameters would help clarify the processes through which reduced food accessibility influences decision-making. Fifth, because the manipulation of food accessibility combined physical effort with cognitive challenges, the relative contribution of these components remains unknown. Future studies should experimentally dissociate energetic expenditure, motor demands, and cognitive complexity. Finally, animals were tested in a probabilistic reversal-learning task. Whether similar benefits generalize to other domains of executive function, such as working memory, inhibitory control, or cognitive flexibility, remains to be determined.

In conclusion, rats required to exert effort to obtain food exhibited improved decision-making as compared with animals kept *ad libitum*, despite comparable task engagement and learning trajectories. These findings provide evidence that food accessibility is an important determinant of cognitive performance and suggest that repeated exposure to biologically relevant learning challenges may enhance the efficiency of reward-guided decision-making. More broadly, the results support the idea that incorporating ethologically meaningful demands into laboratory environments may improve both animal welfare and the translational relevance of behavioral neuroscience models.

## Authors contribution

J.L.P.V. conceived the study. J.L.P.V., M.A.B. and C.I.G.M. designed the study. J.L.P.V., V.H.N., F.J.A., M.A.B., C.I.G.M. and M.B.G. developed the methodology. C.I.G.M., M.A.B. and V.H.N. collected the data. C.I.G.M. and M.A.B. performed the statistical analyses and C.I.G.M. and J.L.P.V. interpreted the results with input from all authors. J.L.P.V. drafted the original manuscript. M.A.B., V.H.N., M.B.G., and C.I.G.M. critically reviewed and revised the manuscript. All authors approved the final version of the manuscript and agree to be accountable for all aspects of the work.

## Acknowledgements

J.L.P.V. was supported by a Ramón y Cajal fellowship (RYC2019-026380-I) and a Young Researchers Grant (PID2019-108250RJ-I00) from the Spanish Ministry of Science and Innovation; by the Excellence Research Projects Programme of the Galician University System (ED431F 2022/20) and the Competitive Reference Research Group Programme of the Xunta de Galicia (ED431C 2022/05); and by a research grant from the Berce Programme of the Universidade da Coruña. F.J.A. was supported by a Margarita Salas postdoctoral fellowship funded by the Spanish Ministry of Universities under the European Union NextGenerationEU programme. M.B.G. was supported by a Deputación da Coruña Research Fellowship (BINV-CS/2024).

## References

Adida, M., Jollant, F., Clark, L., Besnier, N., Guillaume, S., Kaladjian, A., Mazzola-Pomietto, P., Jeanningros, R., Goodwin, G. M., Azorin, J.-M., & Courtet, P. (2011). Trait-Related Decision-Making Impairment in the Three Phases of Bipolar Disorder. Biological Psychiatry, 70(4), 357–365. 10.1016/j.biopsych.2011.01.018

Alejandro, R. J., & Holroyd, C. B. (2024). Hierarchical control over foraging behavior by anterior cingulate cortex. Neuroscience & Biobehavioral Reviews, 160, 105623. 10.1016/j.neubiorev.2024.105623

Ariesen, A.-M. D., Neubert, J. H., Gaastra, G. F., Tucha, O., & Koerts, J. (2023). Risky Decision-Making in Adults with Alcohol Use Disorder—A Systematic and Meta-Analytic Review. Journal of Clinical Medicine, 12(8), 2943. 10.3390/jcm12082943

Bates, D., Mächler, M., Bolker, B., & Walker, S. (2015). Fitting Linear Mixed-Effects Models Using **lme4**. Journal of Statistical Software, 67(1). 10.18637/jss.v067.i01

Brooks, M., E., Kristensen, K., Benthem, K., J. ,van, Magnusson, A., Berg, C., W., Nielsen, A., Skaug, H., J., Mächler, M., & Bolker, B., M. (2017). glmmTMB Balances Speed and Flexibility Among Packages for Zero-inflated Generalized Linear Mixed Modeling. The R Journal, 9(2), 378. 10.32614/RJ-2017-066

Chen, S., Yang, P., Chen, T., Su, H., Jiang, H., & Zhao, M. (2020). Risky decision-making in individuals with substance use disorder: A meta-analysis and meta-regression review. Psychopharmacology, 237(7), 1893–1908. 10.1007/s00213-020-05506-y

Ching, T.-T., & Hsu, A.-L. (2025). The impacts of different dietary restriction regimens on aging and longevity: From yeast to humans. Journal of Biomedical Science, 32(1), 91. 10.1186/s12929-025-01188-w

Colautti, L., Iannello, P., Silveri, M. C., & Antonietti, A. (2021). Decision making in Parkinson’s disease: An analysis of the studies using the Iowa Gambling Task. European Journal of Neuroscience, 54(10), 7513–7549. 10.1111/ejn.15497

Domínguez-Oliva, A., Hernández-Avalos, I., Bueno-Nava, A., Chávez, C., Verduzco-Mendoza, A., Olmos-Hernández, A., Villanueva-García, D., Avila-Luna, A., Mora-Medina, P., Martínez-Burnes, J., Gálvez-Rosas, A., & Mota-Rojas, D. (2025). Environmental enrichment for laboratory rats and mice: Endocrine, physiological, and behavioral benefits of meeting rodents’ biological needs. Frontiers in Veterinary Science, 12, 1622417. 10.3389/fvets.2025.1622417

Domínguez-Oliva, A., Hernández-Avalos, I., Bueno-Nava, A., Chávez, C., Verduzco-Mendoza, A., Villanueva-García, D., Avila-Luna, A., Olmos-Hernández, A., Martínez-Burnes, J., Gálvez-Rosas, A., & Mota-Rojas, D. (2026). Influence of environmental enrichment on rodents’ brains: Neurochemical and neuroanatomical aspects. Frontiers in Veterinary Science, 13, 1767287. 10.3389/fvets.2026.1767287

Feige-Diller, J., Krakenberg, V., Bierbaum, L., Seifert, L., Palme, R., Kaiser, S., Sachser, N., & Richter, S. H. (2020). The Effects of Different Feeding Routines on Welfare in Laboratory Mice. Frontiers in Veterinary Science, 6, 479. 10.3389/fvets.2019.00479

Ferreira, V. H. B., Simoni, A., Germain, K., Leterrier, C., Lansade, L., Collin, A., Mignon-Grasteau, S., Le Bihan-Duval, E., Guettier, E., Leruste, H., Calandreau, L., & Guesdon, V. (2021). Working for food is related to range use in free-range broiler chickens. Scientific Reports, 11(1), 6253. 10.1038/s41598-021-85867-2

François, M., Fernández-Gayol, O., & Zeltser, L. M. (2022). A Framework for Developing Translationally Relevant Animal Models of Stress-Induced Changes in Eating Behavior. Biological Psychiatry, 91(10), 888–897. 10.1016/j.biopsych.2021.06.020

Frederick, M. J., & Cocuzzo, S. E. (2017). Contrafreeloading in Rats Is Adaptive and Flexible: Support for an Animal Model of Compulsive Checking. Evolutionary Psychology, 15(4), 1474704917735937. 10.1177/1474704917735937

Gershman, S. J., & Uchida, N. (2019). Believing in dopamine. Nature Reviews Neuroscience, 20(11), 703–714. 10.1038/s41583-019-0220-7

Graham, A. L. (2021). Naturalizing mouse models for immunology. Nature Immunology, 22(2), 111–117. 10.1038/s41590-020-00857-2

Green, C. L., Lamming, D. W., & Fontana, L. (2022). Molecular mechanisms of dietary restriction promoting health and longevity. Nature Reviews Molecular Cell Biology, 23(1), 56–73. 10.1038/s41580-021-00411-4

Gu, Y., Zhou, C., Yang, J., Zhang, Q., Zhu, G., Sun, L., Ge, M., & Wang, Y. (2020). A transdiagnostic comparison of affective decision-making in patients with schizophrenia, major depressive disorder, or bipolar disorder. PsyCh Journal, 9(2), 199–209. 10.1002/pchj.351

Hage, P., Jang, I. K., Looi, V., Fakharian, M. A., Orozco, S. P., Pi, J. S., Sedaghat-Nejad, E., & Shadmehr, R. (2023). Effort cost of harvest affects decisions and movement vigor of marmosets during foraging. eLife, 12, RP87238. 10.7554/eLife.87238

Harrison, X. A. (2015). A comparison of observation-level random effect and Beta-Binomial models for modelling overdispersion in Binomial data in ecology & evolution. PeerJ, 3, e1114. 10.7717/peerj.1114

Hauber, W. (2026). Behavioral, neural and neurochemical basis of effort-related decision making. Reviews in the Neurosciences. 10.1515/revneuro-2026-0030

Hill, J. O., & Peters, J. C. (1998). Environmental Contributions to the Obesity Epidemic. Science, 280(5368), 1371–1374. 10.1126/science.280.5368.1371

Hinde, K. J., Benninghoff, A. D., Cho, C. E., & Ward, R. E. (2018). Modeling the Western Diet for Preclinical Investigations. Advances in Nutrition, 9(3), 263–271. 10.1093/advances/nmy002

Hobbiesiefken, U., Mieske, P., Lewejohann, L., & Diederich, K. (2021). Evaluation of different types of enrichment—Their usage and effect on home cage behavior in female mice. PLOS ONE, 16(12), e0261876. 10.1371/journal.pone.0261876

Karaoglan, M., & Karaoglan, M. (2024). The evolution of obesity and the origin of adipose tissue. Obesity Medicine, 52, 100561. 10.1016/j.obmed.2024.100561

Keenan, K. P., Laroque, P., Soper, K. A., Morrissey, R. E., & Dixit, R. (1996). The effects of overfeeding and moderate dietary restriction on Sprague-Dawley rat survival, pathology, carcinogenicity, and the toxicity of pharmaceutical agents. Experimental and Toxicologic Pathology, 48(2-3), 139–144. 10.1016/S0940-2993(96)80034-0

Kentner, A. C., Speno, A. V., Doucette, J., & Roderick, R. C. (2021). The Contribution of Environmental Enrichment to Phenotypic Variation in Mice and Rats. Eneuro, 8(2), ENEURO.0539-20.2021. 10.1523/ENEURO.0539-20.2021

Kuhla, A., Lange, S., Holzmann, C., Maass, F., Petersen, J., Vollmar, B., & Wree, A. (2013). Lifelong Caloric Restriction Increases Working Memory in Mice. PLoS ONE, 8(7), e68778. 10.1371/journal.pone.0068778

Kuznetsova, A., Brockhoff, P. B., & Christensen, R. H. B. (2017). **lmerTest** Package: Tests in Linear Mixed Effects Models. Journal of Statistical Software, 82(13). 10.18637/jss.v082.i13

Lenth, R. V., & Piaskowski, J. (2026). ra. https://CRAN.R-project.org/package=emmeans

Lobo, F., Haase, J., & Brandhorst, S. (2022). The Effects of Dietary Interventions on Brain Aging and Neurological Diseases. Nutrients, 14(23), 5086. 10.3390/nu14235086

Lü, W., Yu, T., & Kuang, W. (2023). Effects of dietary restriction on cognitive function: A systematic review and meta-analysis. Nutritional Neuroscience, 26(6), 540–550. 10.1080/1028415X.2022.2068876

Ma, L., Wang, R., Dong, W., & Zhao, Z. (2018). Caloric restriction can improve learning and memory in C57/BL mice probably via regulation of the AMPK signaling pathway. Experimental Gerontology, 102, 28–35. 10.1016/j.exger.2017.11.013

Maric, I., Krieger, J.-P., Van Der Velden, P., Börchers, S., Asker, M., Vujicic, M., Wernstedt Asterholm, I., & Skibicka, K. P. (2022). Sex and Species Differences in the Development of Diet-Induced Obesity and Metabolic Disturbances in Rodents. Frontiers in Nutrition, 9, 828522. 10.3389/fnut.2022.828522

Martin, B., Ji, S., Maudsley, S., & Mattson, M. P. (2010). “Control” laboratory rodents are metabolically morbid: Why it matters. Proceedings of the National Academy of Sciences, 107(14), 6127–6133. 10.1073/pnas.0912955107

Mattison, J. A., Colman, R. J., Beasley, T. M., Allison, D. B., Kemnid, J. W., Roth, G. S., Ingram, D. K., Weindruch, R., De Cabo, R., & Anderson, R. M. (2017). Caloric restriction improves health and survival of rhesus monkeys. Nature Communications, 8(1), 14063. 10.1038/ncomms14063

Mohebi, A., Wei, W., Pelattini, L., Kim, K., & Berke, J. D. (2024). Dopamine transients follow a striatal gradient of reward time horizons. Nature Neuroscience, 27(4), 737–746. 10.1038/s41593-023-01566-3

Morelli, M., Casagrande, M., & Forte, G. (2022). Decision Making: A Theoretical Review. Integrative Psychological and Behavioral Science, 56(3), 609–629. 10.1007/s12124-021-09669-x

Nakagawa, S., Lagisz, M., Hector, K. L., & Spencer, H. G. (2012). Comparative and meta-analytic insights into life extension via dietary restriction. Aging Cell, 11(3), 401–409. 10.1111/j.1474-9726.2012.00798.x

Oraha, J., Enriquez, R. F., Herzog, H., & Lee, N. J. (2022). Sex-specific changes in metabolism during the transition from chow to high-fat diet feeding are abolished in response to dieting in C57BL/6J mice. International Journal of Obesity, 46(10), 1749–1758. 10.1038/s41366-022-01174-4

Pani, G. (2015). Neuroprotective effects of dietary restriction: Evidence and mechanisms. Seminars in Cell & Developmental Biology, 40, 106–114. 10.1016/j.semcdb.2015.03.004

Perugini, A., Ditterich, J., Shaikh, A. G., Knowlton, B. J., & Basso, M. A. (2018). Paradoxical Decision-Making: A Framework for Understanding Cognition in Parkinson’s Disease. Trends in Neurosciences, 41(8), 512–525. 10.1016/j.tins.2018.04.006

Puppe, B., Ernst, K., Schön, P. C., & Manteuffel, G. (2007). Cognitive enrichment affects behavioural reactivity in domestic pigs. Applied Animal Behaviour Science, 105(1-3), 75–86. 10.1016/j.applanim.2006.05.016

R Core Team. (2023). *R: a language and environment for statistical computing* [Manual]. R Foundation for Statistical Computing. https://www.R-project.org/

Rahman, S., Sahakian, B. J., Cardinal, R. N., Rogers, R. D., & Robbins, T. W. (2001). Decision making and neuropsychiatry. Trends in Cognitive Sciences, 5(6), 271–277. 10.1016/S1364-6613(00)01650-8

Richmond, A. A. H., Heinen, V. K., Welklin, J. F., Sonnenberg, B. R., Haley, S. M., Whitenack, L. E., Bridge, E. S., & Pravosudov, V. V. (2026). More experience in the initial learning of spatial information improves cognitive flexibility in food-caching chickadees. Animal Cognition, 29(1), 8. 10.1007/s10071-025-02024-2

Rosenberger, K., Simmler, M., Nawroth, C., Langbein, J., & Keil, N. (2020). Goats work for food in a contrafreeloading task. Scientific Reports, 10(1), 22336. 10.1038/s41598-020-78931-w

Rothkoff, L., Feng, L., & Byosiere, S.-E. (2024). Domestic pet dogs (Canis lupus familiaris) do not show a preference to contrafreeload, but are willing. Scientific Reports, 14(1), 1314. 10.1038/s41598-024-51663-x

Singh, R., Lakhanpal, D., Kumar, S., Sharma, S., Kataria, H., Kaur, M., & Kaur, G. (2012). Late-onset intermittent fasting dietary restriction as a potential intervention to retard age-associated brain function impairments in male rats. AGE, 34(4), 917–933. 10.1007/s11357-011-9289-2

Speakman, J. R. (2013). Evolutionary Perspectives on the Obesity Epidemic: Adaptive, Maladaptive, and Neutral Viewpoints. Annual Review of Nutrition, 33(1), 289–317. 10.1146/annurev-nutr-071811-150711

Speakman, J. R. (2020). Why does caloric restriction increase life and healthspan? The ‘clean cupboards’ hypothesis. National Science Review, 7(7), 1153–1156. 10.1093/nsr/nwaa078

Speakman, J. R., & Elmquist, J. K. (2022). Obesity: An evolutionary context. Life Metabolism, 1(1), 10–24. 10.1093/lifemeta/loac002

Sutton, R. S., & Barto, A. G. (2018). Reinforcement learning: An introduction (Second edition). The MIT Press.

Verdejo-García, A., Alcázar-Córcoles, M. A., & Albein-Urios, N. (2019). Neuropsychological Interventions for Decision-Making in Addiction: A Systematic Review. Neuropsychology Review, 29(1), 79–92. 10.1007/s11065-018-9384-6

Winstanley, C. A., & Floresco, S. B. (2016). Deciphering Decision Making: Variation in Animal Models of Effort- and Uncertainty-Based Choice Reveals Distinct Neural Circuitries Underlying Core Cognitive Processes. The Journal of Neuroscience, 36(48), 12069–12079. 10.1523/JNEUROSCI.1713-16.2016

Wispinski, N. J., Gallivan, J. P., & Chapman, C. S. (2020). Models, movements, and minds: Bridging the gap between decision making and action. Annals of the New York Academy of Sciences, 1464(1), 30–51. 10.1111/nyas.13973

Witte, A. V., Fobker, M., Gellner, R., Knecht, S., & Flöel, A. (2009). Caloric restriction improves memory in elderly humans. Proceedings of the National Academy of Sciences, 106(4), 1255–1260. 10.1073/pnas.0808587106

Zentall, T. R. (2021). Effect of Environmental Enrichment on the Brain and on Learning and Cognition by Animals. Animals, 11(4), 973. 10.3390/ani11040973

